# Soil and water microbial communities: A source of novel antibiotics for sustainable plant disease control

**DOI:** 10.1101/2025.02.06.636970

**Authors:** Navid Iqbal, Noreen Asim, Shahana Malik, Sartaj Khan, Sadia Akbar, Youngchul Yoo, Irfan Ullah, Luo liu, Shahin Shah Khan, Sang Won Lee

## Abstract

Pathogenic microorganisms pose a significant threat to plant health and agricultural productivity worldwide. Herein, this study explores the potential of natural microbial communities as sources of biopesticides. We analyzed 20 soil and water samples from remote areas, isolating 121 colonies using nutrient agar. Antibacterial activity was assessed through agar diffusion, revealing that 12 colonies exhibited significant inhibitory effects against various pathogenic bacteria. Measurement of inhibition zones indicated that these colonies could produce antibiotics. Subsequently, molecular characterization with 16S rRNA and BLAST searches of 12 unidentified sequences highlighted substantial microbial diversity within the samples. This research underscores the promise of these natural microbial communities in developing biocontrol agents for managing plant diseases in greenhouse and field environments. Extraction and purification techniques could further refine these antibiotics, advancing sustainable agricultural practices.

## Introduction

Plant diseases cause substantial reduction in crop growth, production, and storage. Like other living organisms, plants are susceptible to many infections and diseases caused by several pathogens, pests, and microbes of different origins^1^. These diseases can cause minor damage to plants such as growth retardation or can be deadly to plant life in the severe case^2-4^. Nowadays, there is an increased demand for safer, sustainable, and ecosystem-friendly strategies^5-7^. As resultantly strict rules and regulations for the commercialization and production of pesticides as well as alternative strategies like genetic resistance of crops to pathogens and uses of biopesticides have also increased^8^.

Microorganisms have been used to combat plant pathogens for many years. The best alternative biological control for plant pathogens is antibacterial fungi and bacteria^9^. Another strategy for plant protection involves pre-inoculating plant growth-promoting bacteria (PGPB) to induce systemic resistance in crop roots^10^. Microorganism-based biopesticide products have a wide range of applications^11^. They are used for disease control in open fields and greenhouses for various crops, including cereals, legumes, fruits, flowers, and ornamental plants that are infected by soil-borne, foliar, or post-harvest pathogens^12^. These beneficial microorganisms are mainly isolated from the underground or aerial parts of plants that are less or less affected by a pathogen than nearby devastated groups of plants of the same species^13^.

Soil is a diverse source for discovering novel antibiotics. It is rich in microorganisms that can produce antibiotics^14^. While most effective antibiotics were discovered in the 20^th^ century from soil microorganisms, they now show reduced effectiveness against multidrug-resistant pathogens^15^. Uncultured microorganisms in the soil are a rich source of new antibiotics^16-19^. Approximately 75 to 80% of beneficial and commercially valuable antibiotics are obtained from the Streptomyces genus, which makes up 50% of soil microbes^20^. Similarly, the water environment is becoming increasingly recognized as a rich and untapped source of beneficial novel microbes^21^. Freshwater alone is believed to contain a large variety of microorganisms with unique biological characteristics that enable them to thrive in challenging environmental conditions, potentially leading to the production of new secondary metabolites^22,23^. Spring water has medicinal uses, and there is a possibility that the bacteria present in it can produce metabolites that combat diseases^24^.

Herein, this research aims to explore microorganisms capable of producing antibiotics and to evaluate their effectiveness against different pathogens by examining their in-vitro antibacterial activity. Soil and water samples were collected from various remote areas of Khyber Pakhtunkhwa, Pakistan such as district Swat, district Buner, district Dir, and district Shangla. These areas were chosen due to the high likelihood of isolating novel antibiotic-producing soil microorganisms.

### Methodology Sample collection

Soil and water samples were collected from districts Swat, Buner, Dir, and Shangla (Table 1). Samples were gathered from remote areas with minimal human activity, increasing the likelihood of discovering new microbes with novel antibiotics. Soil samples were collected using a trowel to remove debris and stored in a sterilized plastic bag^25^. The water samples were collected in 500 mL sterilized, airtight bottles, using a sterile spatula for residue collection^26^.

**Table 1.** Sources and locations of the isolated colonies.

| Isolates | Sources | District | latitude | longitude |
| --- | --- | --- | --- | --- |
| Isolate 3 | Soil | Swat | 35.4902° N | 72.5796° E |
| Isolate 4 | Soil | Swat | 35.4902° N | 72.5796° E |
| Isolate 5 | Water | Dir | 34.8718° N | 71.8753° E |
| Isolate 8 | Soil | Swat | 35.4902° N | 72.5796° E |
| Isolate 9 | Soil | Swat | 34.7526° N | 72.2986° E |
| Isolate 10 | Water | Swat | 35.4902° N | 72.5796° E |
| Isolate 12 | Water | Swat | 35.4902° N | 72.5796° E |
| Isolate 14 | Water | Buner | 34.3943° N | 72.6151° E |
| Isolate 15 | Soil | Swat | 35.1404° N | 72.5353° E |
| Isolate 18 | Water | Swat | 35.4902° N | 72.5796° E |
| Isolate 19 | Soil | Swat | 34.7526° N | 72.2986° E |
| Isolate 20 | Soil | Swat | 35.4902° N | 72.5796° E |

### Isolation of bacteria from soil and water

Water and soil samples were serially diluted for bacterial isolation. In brief, 1 mL of water was diluted in 9 mL of distilled water, while 1 gram of soil was dissolved in a NaCl solution. The resulting dilutions were then spread onto nutrient agar plates. Distinct bacterial colonies were selected, and pure cultures were obtained. These isolated colonies from the soil and water samples were subsequently cultured in nutrient broth (NB) to evaluate their antibacterial activity^27-29^ (**Table 1)**.

### Pathogenic strains

Six pathogenic strains acquired from the Department of Plant Pathology, The University of Agriculture, Peshawar, were used as test organisms: *Xanthomonas oryzae pv orayzae (Xoo), Erwinia carotovora pv carotovora (Ecc), Pseudomonas syringae (P. syringae), Xanthomonas campestris (X. campestris), Xanthomonas citria (X. citria)*, and *Ralstonia solanacearum (R. solanacearum)*.

### Antibacterial activity

Antibacterial activity against pathogenic bacteria, *i*.*e*., *Xoo, Ecc, P. syringae, X. campestris, X. citria*, and *R. solanacearum*, was assessed using agar diffusion method. All the isolated bacterial strains were cultured in nutrient broth for 24 hours at 37°C. Each bacterial isolates were centrifuged and proceed to grow overnight for further characterization. Further, all the pathogenic strains were spreads uniformly on the plates and paper disc were immersed in supernatants of the isolates for 30 minuets and were placed on the agar plates inoculated with pathogenic strains. After 24 hours incubation, antibacterial efficacy was evaluated by measuring the diameter of the zones of inhibition^30,31^.

### Characterization of isolated bacterial strains

The morphological features of isolates demonstrating antibacterial activity were observed using various biochemical tests and molecular characterization was conducted using 16S rRNA sequencing^32-35^. DNA extraction was performed using the phase lock gel method^36^. The DNA was amplified using 16S rRNA primers. The forward primer (FD1) sequence was AGAGTTTGATCCTGGCTCAG and the reverse primer (rP1) sequence was ACGG(ACT)TACCTTGTTACGACTT. The PCR reactions were conducted and DNA fragments were amplified to approximately 1500 bp. The PCR product was commercially sequenced (Macrogen, Korea).

### Phylogenetic analysis

The sequences were modified, and aligned with ClustalX 2.1 and MEGA 11 software before being compared to GenBank entries using the Basic Local Alignment Search Tool (BLAST) to establish their resemblance to closely related taxa. Gene sequences were retrieved from NCBI and organized in Notepad in a tabular format, listing gene IDs, descriptions, and ancestry information. ClustalX 2.1, an offline alignment tool, was used to evaluate the percentage of similarity and identity among the sequences. Phylogenetic analysis was conducted with MEGA 11 software, constructing a tree that mapped the evolutionary history of the genes.

### Statistical Analysis

All experiments were performed in triplicate, and the reported data are averages. Microsoft Excel was used to calculate means and standard deviation.

## Results

### Isolation and morphological assessment of bacteria

A total of 20 samples were collected from various remote areas in Khyber Pakhtunkhwa, including districts Swat, Buner, and Dir. 12 isolates are shown in Table 1 and Serial dilutions were performed, followed by spreading the samples on nutrient agar media. The plates were left to incubate for three days at a temperature of 28°C. On plates, distant colonies were observed and pure colonies were obtained by streaking them on nutrient agar media. The incubation conditions remained consistent. A total of 121 distinct colonies were identified based on their morphological characteristics, including size, shape, texture, margins, and elevation (**Figure S1**). The isolated colonies were further characterized using biochemical tests and molecular analysis.

### Antibacterial activity

The bacterial isolates were evaluated for their antagonistic ability against various test microbes, including *Xoo, Ecc, P. syringae, X. campestris, X. citria, R. solanacearum*, on nutrient agar media. A 100 µL volume of each bacterial isolate was spot-inoculated at designated positions on the plates. Following inoculation, the plates were incubated for 24 hours to assess the antagonistic potential of the isolates against the pathogenic strains (Figure S2).

The results showed that several isolates effectively inhibited the growth of *Xoo*, indicating their potential as biocontrol agents Among them, isolates 5, 10, 12, and 14 emerged as promising candidates for further investigation. Isolate 12 showed the highest antibacterial activity, producing the largest zone of inhibition, measuring approximately 2.83 cm. Isolates 8, 9, 15, 18, 19, and 20 exhibited moderate antibacterial activity, with inhibition zones ranging between 1 to 2 cm (Figure 1A). Against *Ecc*, which causes soft rot in various crops, isolate 4 showed the most potent antibacterial action, with a zone of inhibition of approximately 2.4 cm. Followed by, isolate 19 exhibited strong antibacterial activity, producing a 2.1 cm inhibition zone. Other isolates, including 8, 9, 10, 12, 18, and 20, displayed moderate activity, with zones of inhibition between 1.2 and 1.6 cm (Figure 1B). Antibacterial activity against *P. syringae* was observed in isolates 3, 9, 10, 18, 19, and 20. Isolate 18 produced the largest zone of inhibition at 3.033 cm, while isolate 10 showed the smallest zone, measuring 0.966 cm (Figure 1C). Out of all the isolated bacteria, eight isolates showed antibacterial activity against *R. solanacearum*. Isolate 19 had the largest zone of inhibition 2.8 cm followed by isolates 10, 18, and 19 were recorded as 2.26, 2.4, and 2.06 cm, respectively. Isolates 3, 8, 9, and 20 exhibited lower activity, with inhibition zones ranging from 1.5 to 1.9 cm (Figure 1D). Regarding activity against *X. campestris*, isolates 8, 9, 10, 12, 14, 15, 18, 19, and 20 exhibited significant antibacterial potential. Isolate 12 demonstrated the highest efficacy, while isolates 8 and 9 showed moderate activity. Isolates 14 and 15 had noticeable, though less pronounced, effects compared to isolate 12, while isolates 18, 19, and 20 exhibited modest but significant activity (Figure 1E). Furthermore. the antibacterial efficacy of the isolates was tested against *X. citri*. Out of 12 isolates, isolates 8, 12, 18, 19, and 20 showed significant antibacterial activity, with isolate 19 being the most effective, followed by isolate 12. The remaining isolates exhibited varying levels of activity, from moderate to low (Figure 1F).

**Figure 1.**
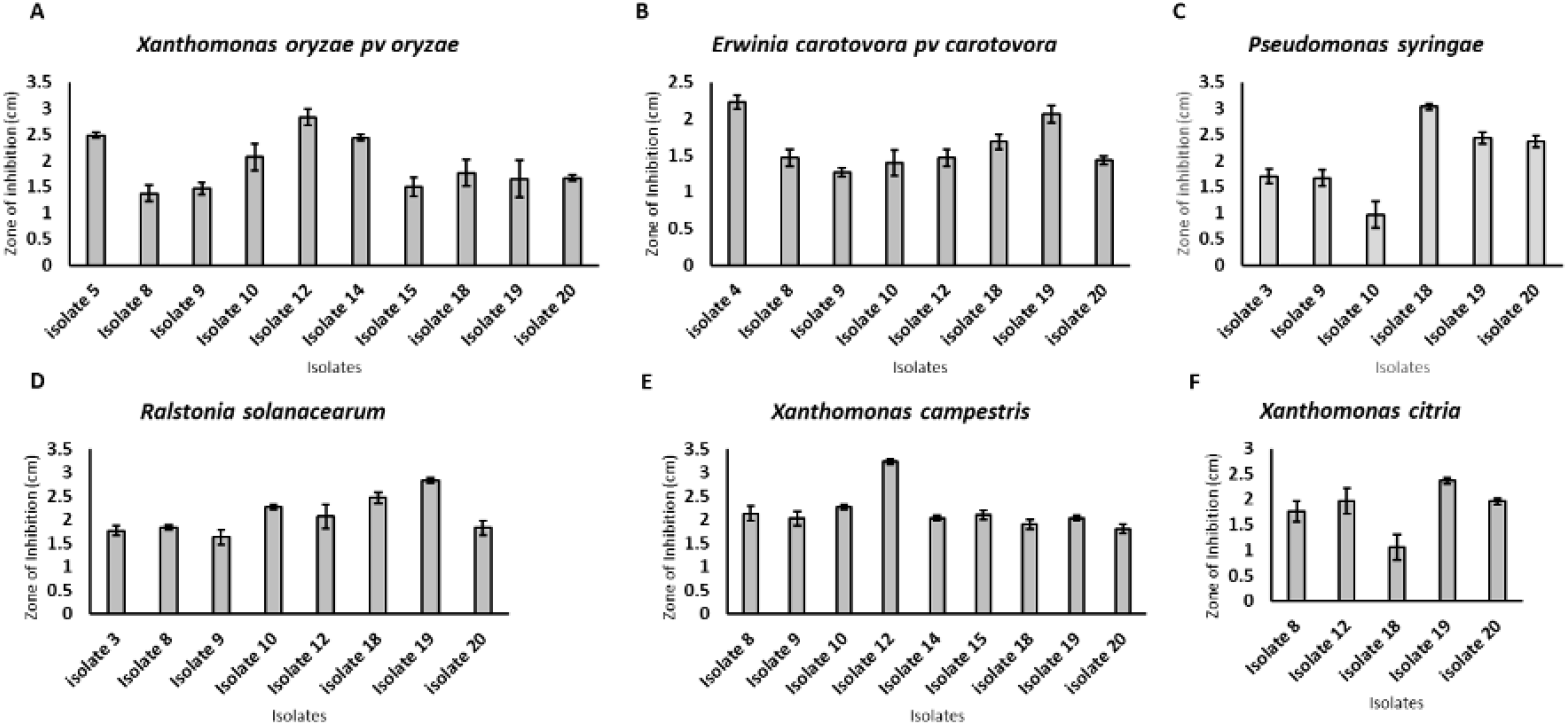
Antibacterial activity of soil and water bacterial isolates against various pathogenic strains: zone of inhibition (cm) measured for different isolates. The data was taken in replicates. (n=3).

In conclusion, isolate 12 consistently exhibited excellent antibacterial activity across multiple pathogens, making it the most promising candidate for further study as a biocontrol agent. Other isolates, such as 19 and 20, showed significant potential against specific pathogens.

### Biochemical characterization of the isolated bacteria

The morphological and biochemical characteristics of the isolated colonies exhibit distinct properties potentially associated with their antibacterial activities. The isolates display diverse colony morphology characterized by a spectrum of colors, including white, yellowish, and dark yellow, alongside various shapes such as irregular, circular, and round. Margins exhibit variations from entire to undulate, elevations fluctuate between flat and raised, and textures can be categorized as smooth or rough. The observed morphological differences may suggest variability in extracellular structures, potentially influencing surface adhesion and environmental resilience, factors that could enhance their capacity to synthesize antibacterial compounds. Rough textures may correlate with heightened extracellular polymer production, facilitating biofilm formation and protection, whereas smooth textures may suggest a streamlined structure beneficial in particular environmental niches.

Biochemical tests reveal significant diversity in metabolic profiles, as demonstrated by Gram staining, catalase, oxidase, urease, and Citrate tests. The Gram staining results indicate the presence of both Gram-positive and Gram-negative bacteria among the isolates, demonstrating diversity in cell wall structures (Table 3). Gram-positive isolates possess thick peptidoglycan layers that may enhance metabolite production, whereas Gram-negative isolates, characterized by an additional outer membrane, may exhibit distinct antimicrobial production or resistance mechanisms. Most isolates exhibit positive catalase activity, indicating an aerobic or facultative anaerobic characteristic, as catalase functions to neutralize hydrogen peroxide, a byproduct of aerobic metabolism. Oxidase results exhibit variability, suggesting differences in electron transport chains among the isolates, potentially linked to variations in energy production pathways and metabolic capabilities. The urease test indicates that numerous isolates possess the ability to hydrolyze urea, resulting in ammonia production that elevates the local pH. The activity of urease may improve survival in acidic environments or facilitate niche adaptation. The TSI test results underscore metabolic diversity, as certain isolates generate gas, others produce hydrogen sulfide (H_2_S), while some exhibit no fermentation. Isolate 3 generates both gas and H_2_S, signifying active fermentation and sulfur metabolism, which may affect its antibacterial properties by establishing an unfavorable environment for competing microbes.

In contrast, isolate 10 exhibits an alkaline slant without gas or H_2_S production, indicating a lack of sugar fermentation and potentially the utilization of alternative metabolic pathways. The presence of black precipitate in certain isolates, indicative of H_2_S production, suggests the ability to reduce sulfur-containing compounds, potentially contributing to their antibacterial efficacy through the generation of reactive sulfur species (Tables 2 and 3).

**Table 2.** Morphological characteristics of isolated colonies that show antibacterial activity.

| **Isolate** | **Isolate** | **Color** | **Shape** | **Margins** | **Elevation** | **Texture** |
| --- | --- | --- | --- | --- | --- | --- |
| **Isolate 3** | 3 | Yellowish | Irregular | Entire | Flat | Rough |
| **Isolate 4** | 4 | White | Rod | Entire | Flat | Smooth |
| **Isolate 5** | 5 | Yellowish | Round | Undulate | Raised | Smooth |
| **Isolate 8** | 8 | Off white | Irregular | Entire | Flat | Rough |
| **Isolate 9** | 9 | White | Circular | Entire | Raised | Smooth |
| **Isolate 10** | 10 | Dark yellow<br>w | Irregular | Undulate | Raised | Rough |
| **Isolate 12** | 12 | Yellowish | Irregular | Entire | Raised | Smooth |
| **Isolate 14** | 14 | Off white | Irregular | Undulate | Raised | Rough |
| **Isolate 15** | 15 | White | Circular | Entire | Raised | Smooth |
| **Isolate 18** | 18 | White | Irregular | Undulate | Flat | Rough |
| **Isolate 19** | 19 | Yellowish | Circular | Undulate | Raised | Smooth |
| **Isolate 20** | 20 | Creamy | Irregular | Undulate | Raised | Rough |

**Table 3.** Morphological and biochemical analysis of the isolated bacterial strains.

| **Isolates** | **Gram staining** | **Appearance** | **Catalase test** | **Oxidase test** | **Urease test** | **Citrate test** | **TSI test** |
| --- | --- | --- | --- | --- | --- | --- | --- |
| **Isolate 3** | - | Rods | + | - | + | - | Red slant, yellow butt, No gas, and H<sub>2</sub>S production |
| **Isolate 4** | + | Rods | + | - | + | - | Yellow slant and red butt, No gas, and H<sub>2</sub>S production |
| **Isolate 5** | + | Rods | + | + | - | - | Red slant, yellow butt, gas, and H<sub>2</sub>S production |
| **Isolate 8** | + | Rods | + | + | + | + | Red slant and butt. No gas production |
| **Isolate 9** | - | Cocci | + | - | + | + | Red slant, yellow butt, No gas, and H<sub>2</sub>S production |
| **Isolate 10** | + | Rods | + | + | - | + | Alkaline slant with no change in the butt, no H<sub>2</sub>S or gas production |
| **Isolate 12** | + | Rods | - | - | + | - | Red slant, yellow butt, No gas but H<sub>2</sub>S production |
| **Isolate 14** | + | Rods | + | + | + | + | - |
| **Isolate 15** | + | Rods | + | + | + | - | Red slant, yellow butt, No gas, and H<sub>2</sub>S production |
| **Isolate 18** | + | Cocci | + | + | - | - | - |
| **Isolate 19** | + | Rods | + | - | + | + | - |
| **Isolate 20** | + | Rods | = | + | + | + | Red Slant, yellow butt, No gas, and H<sub>2</sub>S production |

### Molecular and phylogenetic analysis of the isolates

DNA was extracted from all isolated colonies. The 16S rRNA gene was amplified using universal primers FD1 and rP1, producing a fragment of approximately 1500 bp. The amplified products were visualized on a 1% agarose gel, with a 1 kb marker was used as a reference (**Figure 2**).

**Figure 2.**
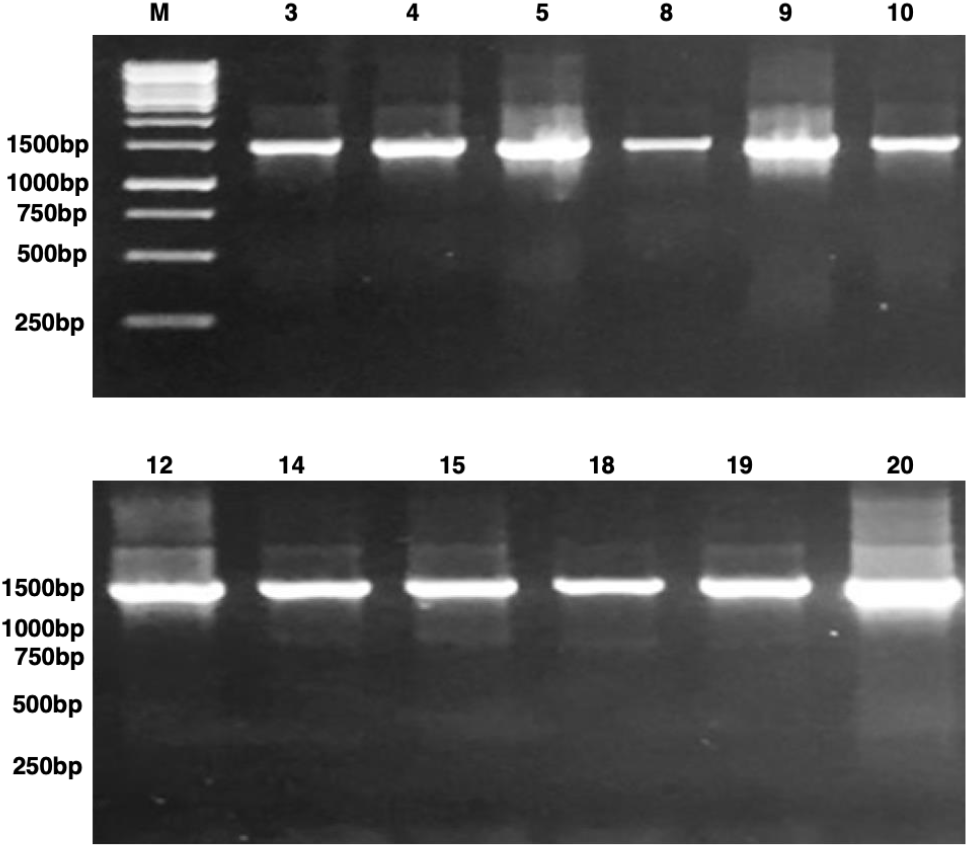
16S rDNA amplified bands of the isolated bacterial strains. M-Marker, and 3, 4, 5, 8, 9, 10, 12, 14, 15, 18, 19, 20 are for Isolates.

The phylogenetic analysis assessed the evolutionary relationships between the bacterial DNA sequences from the isolates and the aligned sequences obtained through BLAST. The study employed the Neighbor-Joining (NJ) algorithm, with evolutionary history and phylogenetic tree construction using MEGA X. The phylogenetic tree offers comprehensive insights into the genetic connections among different bacterial isolates and established bacterial species, with a particular emphasis on the *Bacillus* genus. Isolate 3 and Isolate 20 are positioned next to *Bacillus pumilus*, sharing a node that exhibits strong bootstrap support (99). This strong connection suggests that both isolates could be closely linked to *Bacillus pumilus*, indicating potential strain-level similarity or a recent common ancestor. Isolate 4 is strongly associated with *Bacillus thuringiensis*, evidenced by a bootstrap value of 100. The tight clustering signifies a significant genetic similarity, indicating that isolate 4 may be a strain of *Bacillus thuringiensis*. Isolate 5 clusters containing *Bacillus safensis* and Bacillus sp. NCCP-0439, indicating genetic similarity to *Bacillus safensis*. This clade is supported by a bootstrap value of 100, indicating a strong evolutionary relationship, suggesting that Isolate 5 may be closely related to or a strain of *Bacillus safensis*. Isolate 8 clusters of *Bacillus amyloliquefaciens*, demonstrating a bootstrap value of 100, indicating a strong genetic relationship. This suggests that Isolate 8 may be a member of the same species as *Bacillus amyloliquefaciens* or a closely related strain. Isolate 9 demonstrated close clustering with Bacillus species, indicating a notable genetic similarity and likely close evolutionary relationship with these Bacillus taxa. The bootstrap value of 100 at this node supports the strong association, suggesting isolate 9 may share a recent common ancestor with these species. Isolate 10 demonstrated a close phylogenetic relationship with *Bacillus* sp., as indicated by a bootstrap value of 100. It is incorporated within a robust clade alongside other *Bacillus* species, signifying its membership in a genetically analogous subgroup within the *Bacillus* genus. Isolate 12 clustered with *Clostridium* sp. and *Clostridium* sp. E129, supported by a bootstrap value of 100. This suggests that Isolate 12 exhibits genetic similarities with multiple *Clostridium* species, indicating a close evolutionary relationship within the *Clostridium* genus. Isolate 14, identified inside a clade alongside *Microbacterium paraoxydans* A9s and an *Microbacterium* sp., exhibits strong bootstrap support (100), indicating its association with the *Microbacterium* genus and a likely close kinship to these species, either as a similar or unique strain. Isolate 15, clustered with *Bacillus horikoshii* and displaying a bootstrap value of 100, indicates a substantial genetic affinity to this species, suggesting a strong evolutionary connection and potential strain-level similarities. Isolate 18 has a significant genetic affinity to *Brevundimonas vesicularis* W4 and a *Brevundimonas* species, evidenced by a bootstrap value of 100, signifying its close relationship with the *Brevundimonas* genus and possible classification within this taxonomic group. A bootstrap value of 100 confirms isolate 19’s high affinity for *Bacillus thuringiensis*, pointing to a possible recent common ancestor and hinting that it could be a related strain. (**Figure 3**).

**Figure 3.**
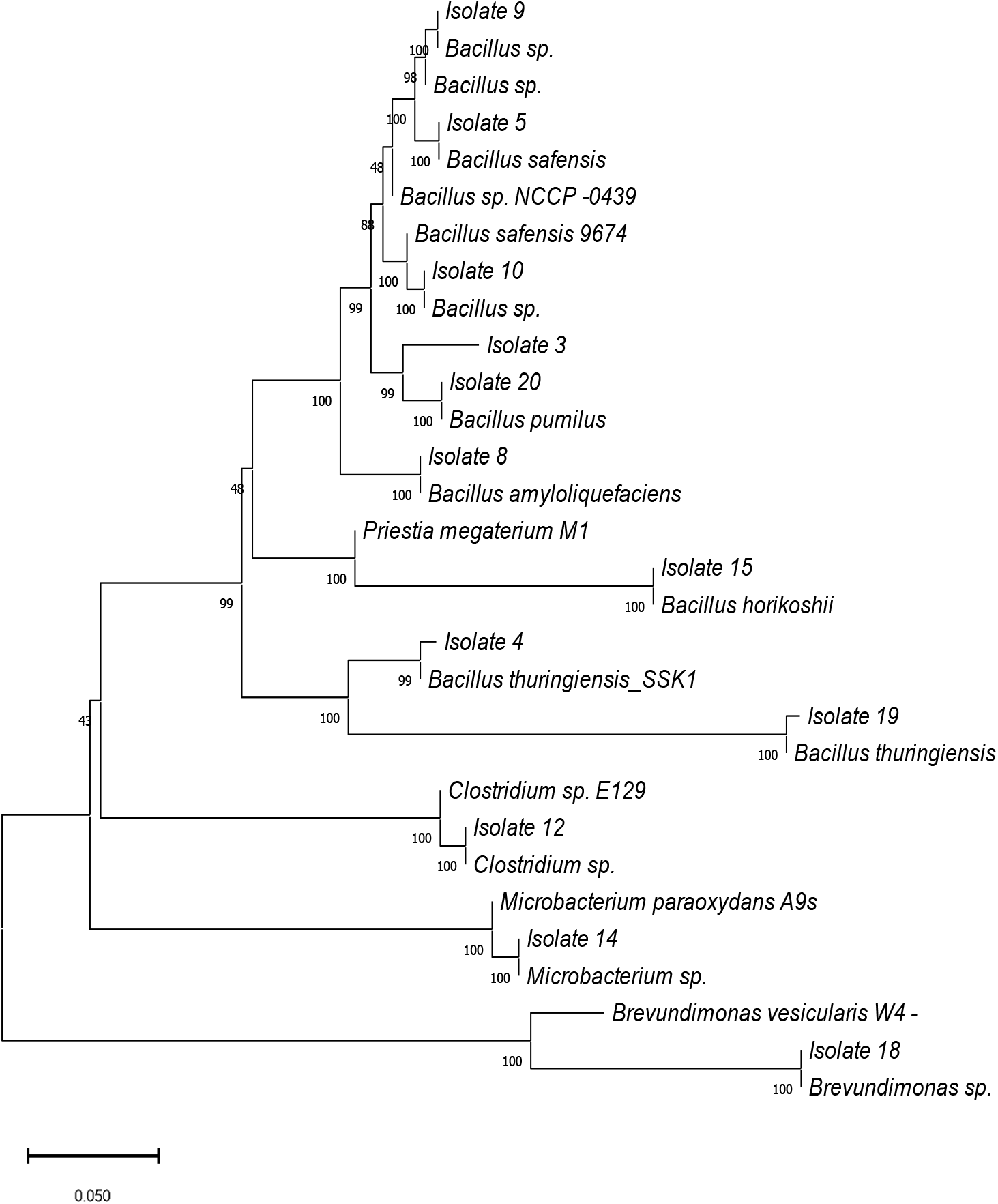
A phylogenetic tree depicting the evolutionary relationships among bacterial isolates, mainly from the genus *Bacillus*, and other species including *Clostridium, Microbacterium*, and *Brevundimonas*. Bootstrap values assigned to each node indicate confidence levels, while the proximity of isolates suggests strain-level similarity. The scale bar indicates genetic distance, illustrating genetic diversity among isolates. This tree demonstrates the genetic diversity and taxonomic relationships among the isolates.

The Sequences from all bacterial isolates were submitted to GenBank for accession numbers. **Table 4** lists the top matching strains, along with their accession numbers and percent identities.

**Table 4.**
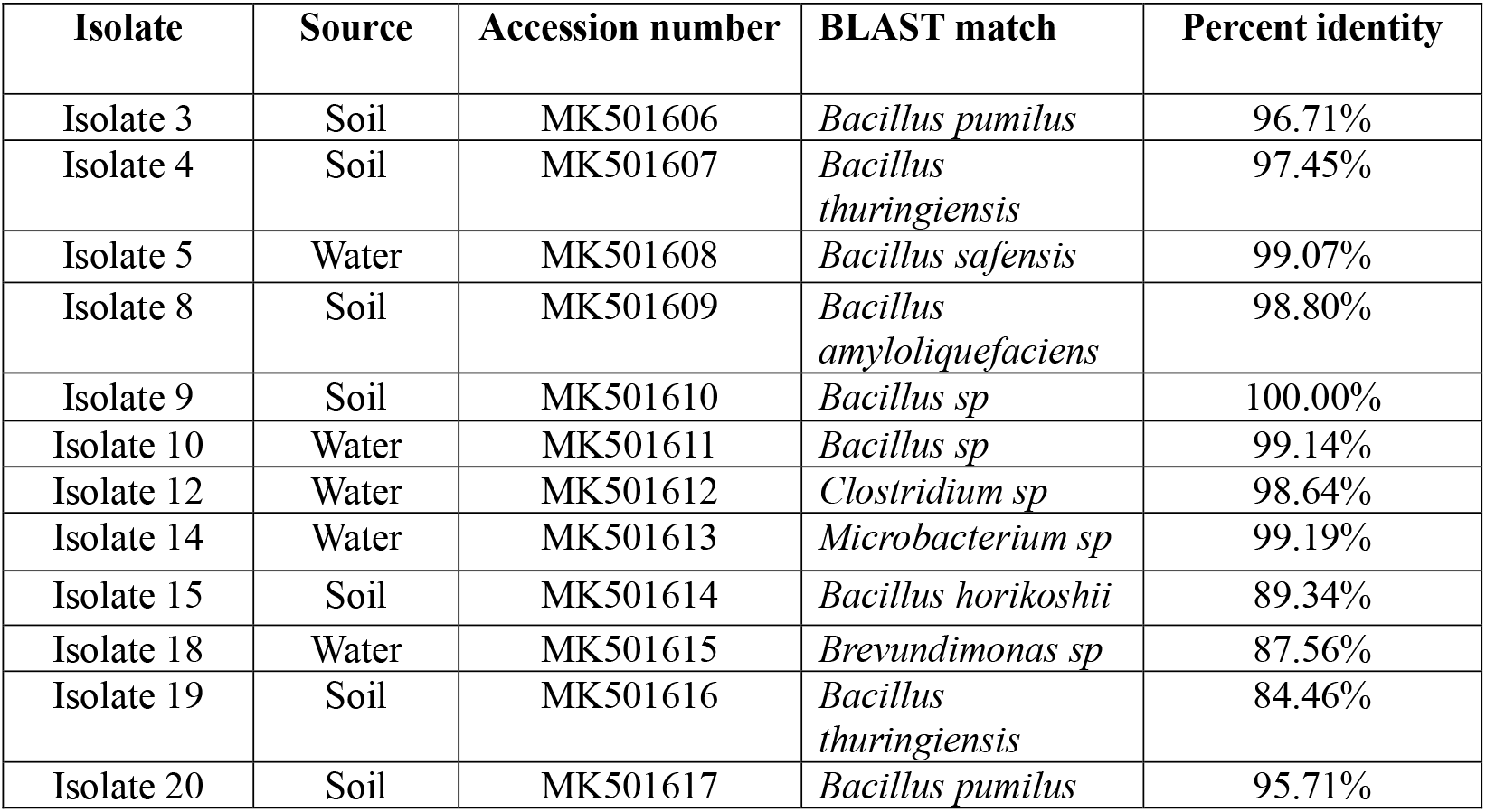
The 16S rDNA sequence analysis of the isolated bacteria.

## Discussion

Continuing the pursuit of new antibiotics from environmental sources is of utmost importance in light of the growing risk posed by antibiotic resistance ^37,38^. Water environments are increasingly being recognized as potential sources of new antimicrobial compounds, alongside soil which has long been known for its antibiotic-producing properties ^38^. Throughout history, the soil has been utilized for its antibiotic properties due to the presence of bacteria, particularly actinomycetes, that are known to produce antibiotics^39^. Further investigation into various ecosystems, along with the use of advanced screening and characterization methods, holds the potential for uncovering novel antibiotics that can effectively combat resistant pathogens. This study investigated the antibacterial properties of bacterial isolates collected from various regions in Khyber Pakhtunkhwa, such as district Swat, Buner, and Dir. A thorough investigation, involving the study of morphological characteristics, biochemical tests, and phylogenetic analysis through 16S rRNA sequencing, provided valuable insights into the potential use of these isolates for plant pathogens.

For screening purposes, a total of twenty samples were collected. These samples were divided equally between water and soil samples, with ten of each. The samples were collected from remote regions of Khyber Pakhtunkhwa, with varying altitudes. A total of 121 colonies were identified based on morphological assessment and examined their antibacterial activity against plant pathogenic strains. The well diffusion assay was utilized to conduct an antibacterial assay. The strains that exhibited a zone of inhibition against pathogenic strains were then analyzed using biochemical and molecular techniques.

Prior studies have demonstrated that soil is an abundant reservoir of actinomycetes, which possess the capacity to synthesize antibiotics that can effectively counteract pathogenic strains. However, the research findings indicated that water sources, in addition to soil, also offered advantages. The majority of the isolates discovered in the Sundarbans grove habitat were of the Streptomyces genus. Among the 54 isolates, only nine exhibited noteworthy antibacterial activity against a diverse array of bacteria and a phytopathogenic fungus^40^. Research has demonstrated that freshwater *pseudomonads* produce chemicals that inhibit the growth of oomycetes, including *Phytophthora sojae* and *Pythium* species. These findings suggest that aquatic isolates could be valuable for developing biocontrol agents^41^. In addition, research using bacterial isolates like *Fluorescent Pseudomonads* and *Trichoderma* spp. has revealed a wide range of antagonistic abilities against *Xanthomonas oryzae pv. oryzae*. Some isolates have shown strong suppression of pathogen development, suggesting their potential for biocontrol^42^. In contrast to these findings, our analysis uncovered a higher percentage of isolates that demonstrated top-notch antibacterial activities. However, the results were not consistent because of the differences in the terrain of the sample site and the specific methodology used. Our research also examined remote locations that have little human interference and no disruptions to the microbial populations. In addition, the collection of samples was place in several different locations, such as the soil around grass near springs, the soil near springs on the mountain’s highest points, and spring water. The soil sample was collected at different depths from 5cm to 20cm. previously, soil samples were collected from various locations including the rhizosphere, cooking house, cattle breeding area, house-waste disposal area, and industrial-waste disposal area. They have collected soil from a depth of up to 11cm^43^. In comparison to their study, our data suggest that there is a shortage or lack of isolates in the sample that was collected from a low depth. On the other hand, a much larger number of isolates were recovered from samples that were taken from significantly deeper regions and from locations that had higher levels of moisture. As a result, it is possible to conclude that bacteria prefer to reside in deeper locations that have an environment that is damp and moist, as opposed to the surface. Furthermore, the antibacterial activity tests revealed that several isolates had substantial inhibitory effects against diverse pathogenic microorganisms. Isolate 12 exhibited the highest level of antibacterial activity against *Xanthomonas oryzae pv. oryzae*, as demonstrated by a zone of inhibition measuring around 2.83 cm or more. These findings align with previous studies that have recognized *Bacillus* species as very proficient producers of antibiotic compounds. The effectiveness of *Bacillus* species, particularly *Bacillus pumilus, Bacillus thuringiensis*, and *Bacillus amyloiquefaciens*, in inhibiting the growth of pathogenic bacteria is widely documented, and this study further supports their potential as agents for biological control ^44-46^. Moreover, the taxonomy of the isolates was well elucidated using biochemical and molecular characterization, which included 16S rRNA sequencing. The classification of the isolates into *Bacillus, Brevundimonas, Rhizobium, Clostridium, Microbacterium*, and uncultured bacterial species highlights the wide variety of bacteria that have antibacterial properties. The majority of the isolates were of *Bacillus* species, which are renowned for their ability to produce a wide range of antimicrobial peptides. The phylogenetic analysis carried out using the Neighbor-Joining method and MEGA 7 software, has uncovered robust evolutionary links between these isolates and well-established antibacterial strains. This research establishes a dependable basis for contemplating their potential application in biocontrol strategies^47-49^. Pakistan has a wide spectrum of biological diversity due to its diverse terrain and geographical dispersion, resulting in a variety of microorganism habitats. Consequently, doing research in this field may be quite beneficial for the purpose of discovering and designing drugs. Several research studies undertaken thus far have been centered on different geographical locations, including the one where our work was conducted. Research was conducted to isolate actinomycetes from several places in Khyber Pakhtunkhwa, Pakistan, including green land, agricultural land, and marshy soil areas^50^. However, during our research, we collected samples from remote locations with limited human interference. Based on the research findings, it is highly likely to detect distinct strains of actinomycetes in various locations in Pakistan. In light of these findings, it is highly recommended to conduct further research to isolate novel bacterial strains at a different location in Pakistan as the sample site. Chemical pesticides have been extensively employed in agriculture for many years; yet, their utilization is accompanied by notable disadvantages. The potential consequences encompass environmental pollution, adverse effects on unintended creatures, emergence of pest resistance, and possible health hazards^51^. Chemical pesticides can have long-lasting effects on the environment, causing harm to the ecosystem by persisting in the soil and water. Not only do they eradicate pests, but they also disrupt the balance of beneficial insects, causing disruptions in natural ecosystems. Insects and other pests can develop resistance to chemical pesticides over time, leading to the need for higher doses or the development of new remedies^52^. As a more sustainable approach to pest management, biopesticides are gaining attention. Biopesticides are made from natural sources like plants, bacteria, fungi, or minerals. Typically, these pesticides are less harmful than traditional ones and have a narrow focus on specific pests, sparing beneficial organisms from harm^51^. Biopesticides often exhibit rapid degradation in the environment, hence minimizing the potential for persistent contamination. They can have a significant impact when used in lesser amounts and are frequently compatible with integrated pest control tactics. Nevertheless, biopesticides may exhibit longer response times in comparison to conventional pesticides and might be more susceptible to variations in environmental circumstances^53^. The current work primarily focuses on the first stage of biopesticide production. We focused on easily accessible natural sources and collected samples without any obstacles. Pakistan possesses abundant natural resources that could harbor unique microorganisms with the ability to produce antibiotics and antifungals. These microorganisms can contribute to the development of a pollution-free environment and agriculture.

Given the limited number of samples and identification methods used in this study, the findings may not offer a complete understanding of the entire identification process with definitive results. Therefore, it is crucial to carry out this study across a wide geographic area, using an advanced identification technique to ensure precise results. To achieve accurate results, it is important to use proper sample techniques. Additionally, including a purification stage with a specified buffer can be beneficial. In conclusion, it is advisable to use natural sources for antibiotic production because of their cost-effectiveness and positive impact on health.

## Conclusions

The study aimed to examine the antibacterial characteristics of the bacterial isolates obtained from several distant regions in Khyber Pakhtunkhwa, including district Swat, Buner, and Dir. Through the use of systematic sampling, serial dilution, and incubation on nutrient agar, a total of 121 bacterial colonies were detected, with each colony being differentiated by its distinct morphological attributes. The isolates were evaluated for their antibacterial activity against different pathogenic strains. The findings revealed that isolates 5, 10, 12, and 14 had good potential for further investigation in biocontrol research. The isolates were classified as members of several genera, such as *Bacillus, Brevundimonas, Clostridium*, and *Microbacterium*,, using molecular characterization and biochemical testing. The findings were validated by phylogenetic analysis, which provided insights into the evolutionary links. This research emphasizes the potential of these isolates as biocontrol agents, offering a natural substitute for artificial pesticides. Additionally, it underscores the significance of conducting additional studies on their practical implementations in the field of agriculture.

## Supporting information

Figure S1. Representative figures of colonies isolated from soil and water

