## Supplementary material for "Soil and water microbial communities: A source of novel antibiotics for sustainable plant disease control": Figure S1. Representative figures of colonies isolated from soil and water

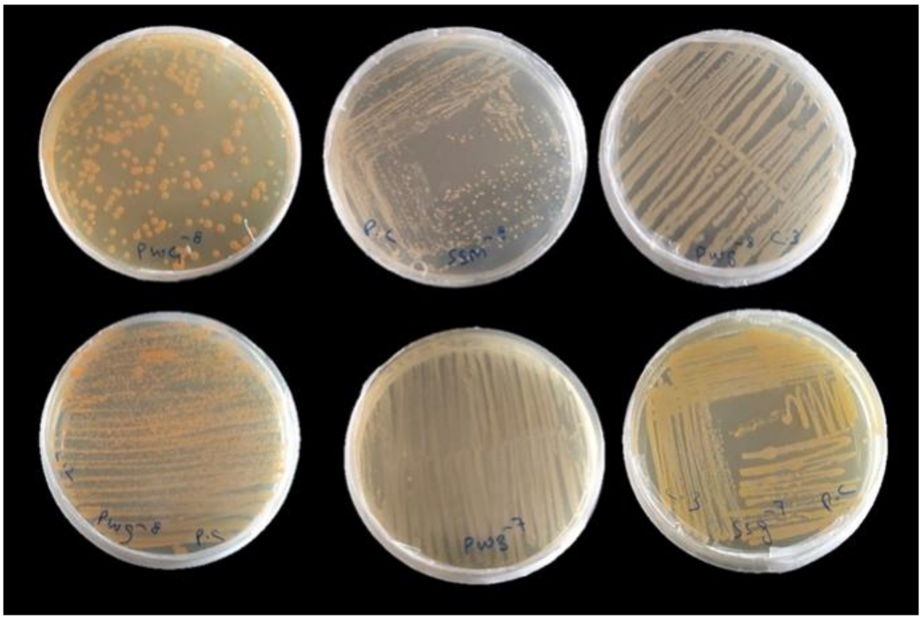

**Figure S1**. Representative figures of colonies isolated from soil and water

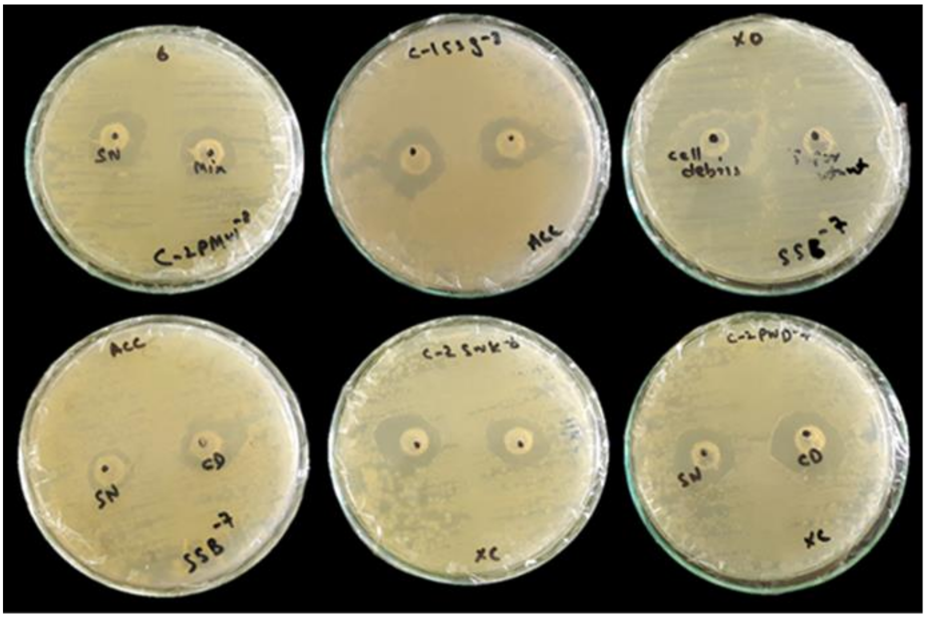


**Figure S2**. Representative figures of antibacterial activity of bacterial strains. Clear zones of inhibition were observed.
